# Assessment of Potential Health and Genetic Impacts in Releasing Confiscated *Paroaria coronata* and *Saltator similis*

**DOI:** 10.1101/2020.03.10.985473

**Authors:** Cláudio E. F. Cruz, Gustavo R. Funkler, André L. S. Zani, Paulo G. C. Wagner, Luciano N. Segura, Inês Andretta, Nelson J. R. Fagundes

## Abstract

Illegal capture and trade of wild birds has long been a threat to biodiversity. Translocation—the release of individuals from one location into another—is a useful conservation tool in the management of species. However, both health (such as different pathogens) and adaptive (such as local adaptation), differences among populations must be taken into account, as both can impact the recipient population negatively. Here, we provide health and genetic information to support release planning for two of the most trafficked Brazilian wild bird species (*Paroaria coronata* and *Saltator similis*). We focused on two fundamental questions: Are there significant differences in pathogen load between wild and captive populations? Is there significant genetic structure among populations? In total, 223 free-living birds were captured, sampled, and released at the same site. Devices and live decoys characteristics were top factors influencing captures. We tested blood, feces, and oropharyngeal swabs from free-ranging (n=101) and confiscated (n=92) birds for Newcastle disease virus, *Salmonella* spp., and *Mycoplasma gallisepticum*. Genetic structure among populations was investigated using mtDNA in a subsample of these birds. We found no evidence for Newcastle disease virus and *Salmonella* spp. in seized and free-living birds from both species. However, seized *P. coronata* and *S. similis* may be potential sources of *M. gallisepticum*. We found significant but low genetic structure among populations occurring in different Biomes (*Φ*_*CT*_=0.26 for *P. coronata*; *Φ*_*CT*_=0.13 for *S. similis*) and no significant structure among populations occurring in the Pampa Biome. These results suggest that while it may be important to screen seized birds for avian pathogens, genetic structure among populations seems to be of lesser concern when planning the release of seized songbirds in the wild.

## Introduction

Illegal trade, along with poaching, habitat loss, and pollution, are top causes of wild bird decline in Brazil and in other developing countries that harbor a huge species diversity [1–4]. The illegal wildlife trade has increased dramatically over the past decade, along with enforcement efforts aimed at mitigating this threat [5]. For confiscated wild birds, despite the shortage of studies on the subject, current translocation guidelines include as an option the euthanasia of individuals belonging to species of low conservation value [6,7]. In part, this policy can be justified on the grounds that a) seized birds may harbor pathogens that will affect the wild population [8] and b) usually, seized birds come from an unknown parental population, and their release in another population may result in outbreeding depression [9,10], resulting, in both cases, in a negative impact on the wild population. Health and genetic studies in wildlife species depend on direct capture for sampling free-ranging individuals and collecting relevant data to evaluate the sanitary and genetic risk of planned translocations. Ideally, such knowledge would allow the translocation of seized and rehabilitated birds to their most probable area of origin, with minimal health risks to the local populations and to the commercial flocks [5–8]. Unfortunately, both genetic and health information on the subject is scarce. Despite the myriad of methods and tools available, mist nets are globally recognized as the top capture method in biological studies [11–13] and are increasingly applied in association with bird lure techniques to improve efficiency, mostly through playback audio [14]. In this sense, the use of artificial [15,16] or live decoys [17], although routinely and widely used in poaching, is only occasionally presented on scientific reports, albeit extensively disseminated in the virtual media.

In this study, we describe the methods for capturing and sampling wild *Paroaria coronata* (Red-crested Cardinal) and *Saltator similis* (Green-winged Saltator)—two of the most heavily trafficked Brazilian wild bird species, which are classified as of least concern on the IUCN red list [18]. Next, we generate sanitary and genetic data to answer two fundamental questions: Are there significant differences in pathogen load between wild and captive populations? Is there significant genetic structure among populations? Finally, we discuss the potential impacts of a translocation program of confiscated conspecifics for both species.

## Materials and Methods

### Ethical statement

This project (no. 23644) was approved by the Ethics Committee on the Use of Animals—UFRGS—and licensed by the *Instituto Chico Mendes de Conservação da Biodiversidade* (ICMBio), under the number 37567. Samples received from Argentina were registered under the export permit code EX-2019-12969382-APN-DNBI#SGP.

### Sampled areas

Sampling areas were selected to include representative areas of the species’ usual distribution, mostly in Rio Grande do Sul state but also in strategic areas from the total species’ distribution (Fig. 1, S1 Table). We captured *S. similis* in two different regions: Pampa and Atlantic Forest. Even though the Pampa is associated with open grasslands [19], we found *S. similis* always associated with forest paths as well as in the Atlantic Forest region. On its turns, we captured *P. coronata* in three different regions: Pampa, Atlantic Forest, and Chaco/Pantanal. In the Atlantic Forest, this species was associated with anthropic deforested paths. The Chaco/Pantanal region is characterized by savannah with a substantial forest component, especially in the Chaco, whose more humid portions are connected to the seasonally flooded Pantanal plain in Brazil [20–22]. Sample size was determined based on both pilot expeditions and on similar studies [23,24]. For the genetic analyses we also included sequences from the GenBank (Bolívia n=1, Boracéia n=1, Corrientes n=2, Mato Grosso n=1) and an additional five *P. coronata* samples were received from the Buenos Aires area, Argentina.

**Fig. 1.**
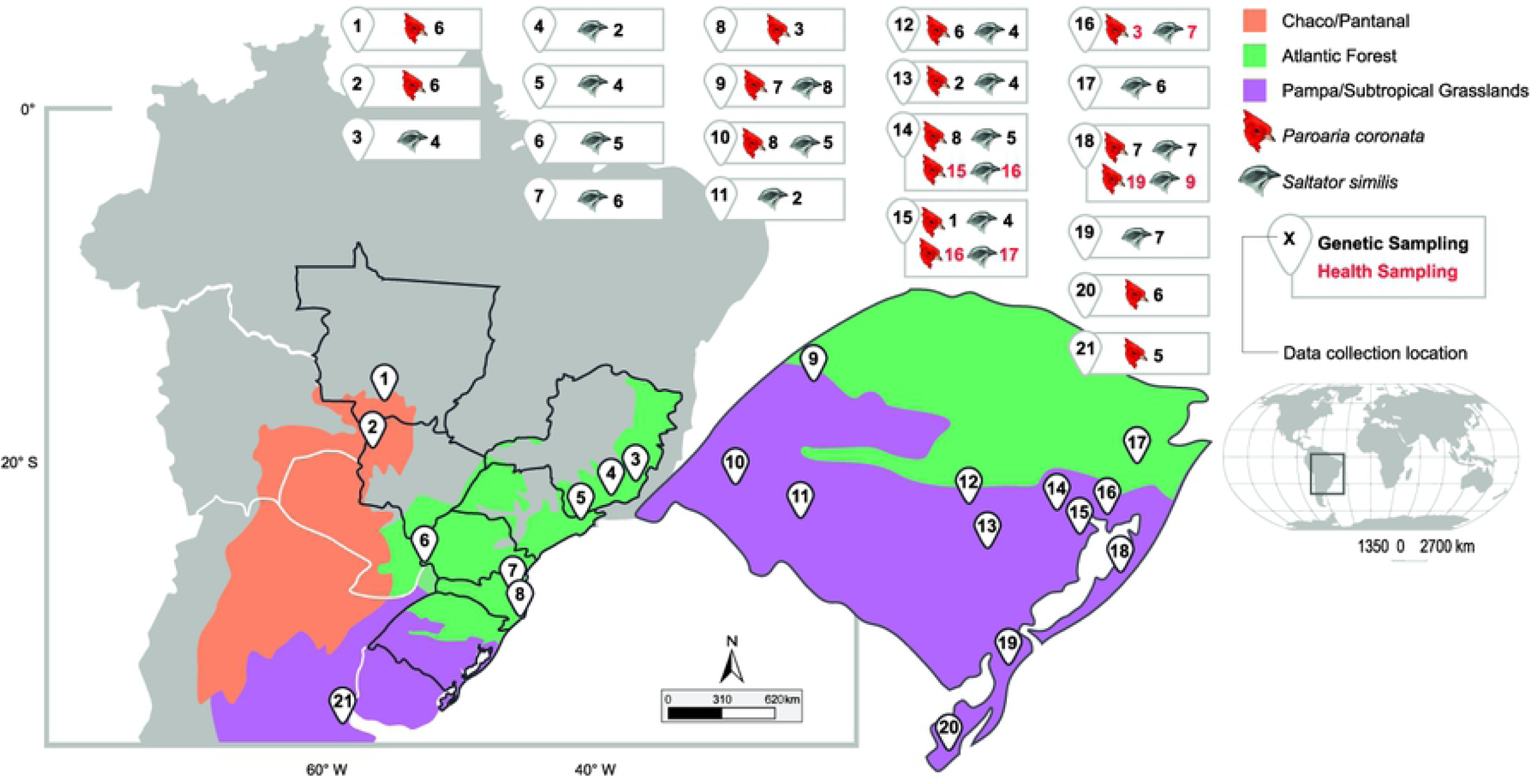
Geographical representation of sampled areas.

### Management of live decoys

Dominant *P. coronata* and *S. similis* of both sexes, identified through bird song defiance (playback) among groups of birds seized from illegal traffic at the *Centro de Triagem de Animais Silvestres* (CETAS/IBAMA), Porto Alegre, RS were tentatively selected, tested, and kept as live decoys for attracting free-ranging conspecific birds. Upon joining the live decoy flock, birds had three consecutive fecal samples negative for parasites. Birds with positive samples were treated (anthelmintic and anticoccidial drugs). Only birds that systematically attracted conspecific birds in fieldwork were kept. Birds that failed 2–4 times consecutively were returned to CETAS. Seed mixtures, commercial rations, fruits, vegetables, arthropods, and minerals composed the birds’ diets. During the capturing time (breeding season—August to February), the birds were kept in cages, and in the rest of the year, they were released in 12–24 m^3^ outdoor aviaries. Bath bowls, sunbath, and cage tray cleaning (sand bedding exchange) were performed every other day.

### Fieldwork

Initially, we used mist nets and cages mounted with net-trapdoors (Fig. 2A) for capturing both species. However, after successive unsuccessful attempts to capture *P. coronata*, a bird trap-loop (Fig. 2B and 2C) was built according to an Indochinese model demonstrated on the internet [25], and used next to the decoy’s cage. All capturing processes were timed.

**Fig. 2.**
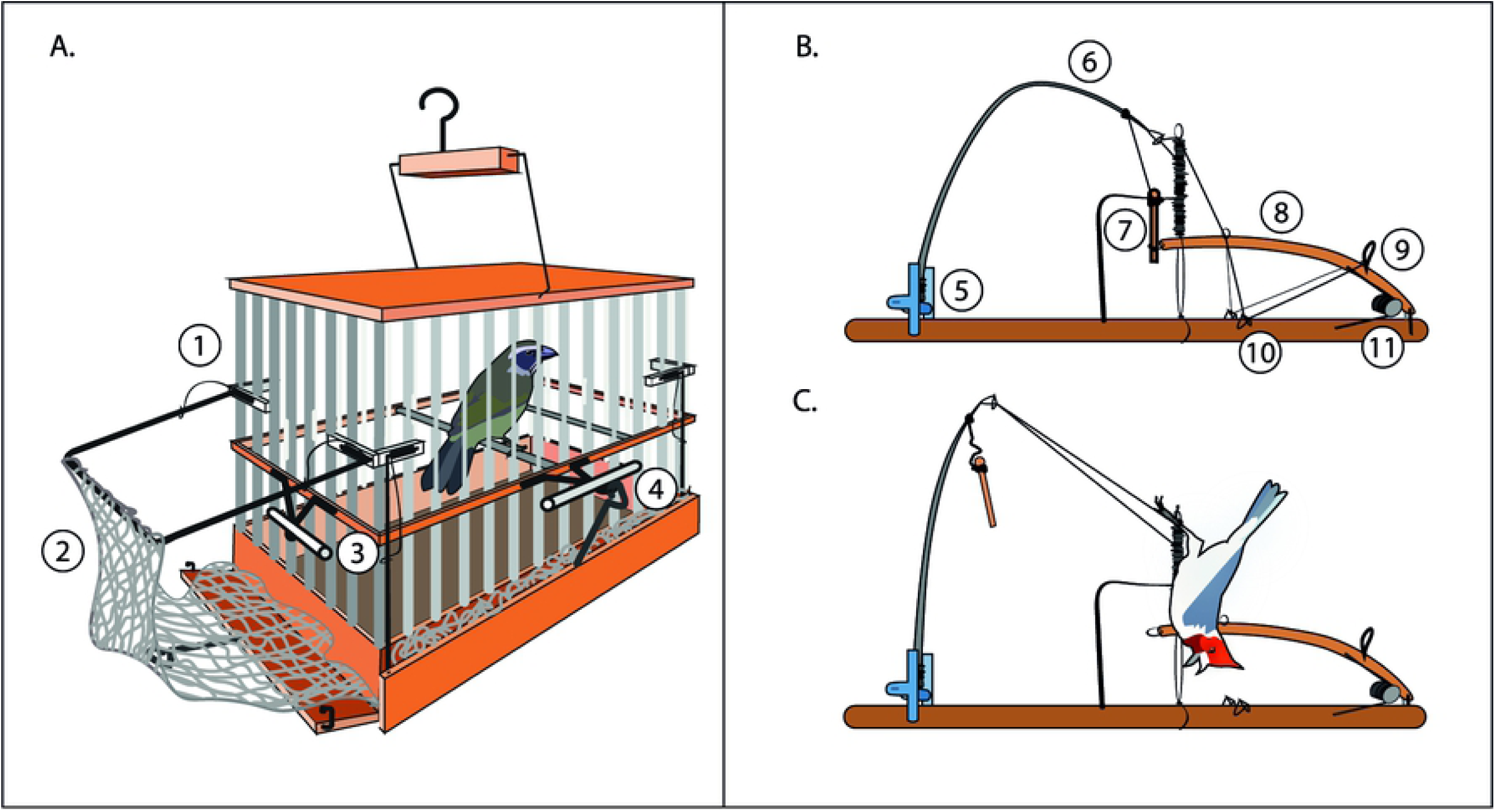
Schematic representation of capturing devices. A: cage with live decoy *Saltator similis*, showing details of spring that closes the net-trapdoor (1), net-trapdoor (2), trigger-perch (3), and metal hook holding the trigger-perch mounted and the net-trapdoor opened (4). B: loop with its parts numbered as the spring that pulls the fishing-rod which pulls the loop (5), fishing-rod (6), pin holding the trigger-perch mounted and fishing-rod (7), trigger-perch (8), brackets (nylon line folds) holding the loop open (9 and 10), and spring holding the trigger-perch up (11). C: *Paroaria coronata* captured in the loop.

Initial sampling was mostly conducted for studying genetic structure in both species. Blood samples (0.1 ml) were obtained by right jugular vein puncture with an insulin syringe [26] and conserved in FTA cards for subsequent genetic applications. A second round of capture expeditions was conducted for obtaining most samples for the sanitary study, in which 0.1 ml of blood (stored in Eppendorf tubes for serum separation), oropharyngeal swab, and feces (mostly after keeping the bird in a holding bag for up to 30 min) were obtained. Such samples were tested for *M. gallisepticum* (MG) and Newcastle disease virus (NDV) serology, MG-PCR, and *Salmonella* spp. isolation, respectively. Samples for the sanitary studies were obtained from populations located up to 200 km from Porto Alegre, to make it possible to deliver them in the laboratory in the same day of collection. Samples were kept under refrigeration. The sampling period included the reproductive seasons of 2017, 2018, and 2019. After sampling, birds were banded and released in the same place of capturing. Additional data is shown in S1 Table.

### Sanitary tests

The microbiological, serological, and molecular tests were performed at the Porto Belo Laboratory, accredited by the Brazilian Ministry of Agriculture, Livestock, and Supply (*Ministério da Agricultura, Pecuária e Abastecimento*—MAPA) to perform official diagnostic tests within the National Plan of Avian Sanity (*Plano Nacional de Sanidade Avícola*—PNSA).

Anti-NDV antibodies were searched for in serum samples by the Hemagglutination Inhibition Test according to the SDA ordinance nº 182, 08 November 1994. *Salmonella* spp. detection followed the methods presented in SDA ordinance nº 126, as of November 03, 1995, after replacing BHI broth by buffered pebble water 1%. Suspected colonies were confirmed and characterized with specific antisera (serum agglutination on blade) at the Fundação Oswaldo Cruz, according to World Health Association protocols [27]. For the detection of anti-MG antibodies, serum samples were tested by a rapid plate agglutination (RPA) test according to SDA ordinance nº 44, as of November 08, 2001. A real-time polymerase chain reaction assay using a Taqman-labeled probe for the detection of *M. gallisepticum* (commercial kit MG—NewGene®) DNA [28] was applied, and the positive results were sent for confirmation with a Multiplex real-time PCR [29] at Simbios Biotechnology.

Serum samples from three *P. coronata* and three *S. similis* vaccinated against MG and NDV served as controls for validation of the MG-qPCR and NDV-HI. Serum samples from these birds were tested before and after vaccination to serve as negative and positive controls, respectively. Vaccination protocols included one ocular drop from live vaccines *M. gallisepticum* (MYCOVAX-TS-11, MERIAL, strain TS-11, P.400/17, V.02/18, November 11, 2017) and NDV (MERIAL, strain La Sota 004/16, ND1873, P.004/16, V.04/18, December 01, 2017), boosters with live vaccines for MG (BIOCAMP, Camp VacMG-F, P.006/2016, V.11/19, February 02, 2018) and NDV (BIOVET, New-Vacin, La Sota, P.007/17, V.05/19, February 23, 2018). An additional booster included an oil inactivated NDV (BIOVET, New-BRONK-VET, P.003/17, V.08/19, virus B1 La Sota, minimum title before inactivation 10^5,3^ DIOE_50_, March 23, 2018). Protocol was based on a previous study on NDV vaccination of wild birds [30]. For the sanitary analyses, we also included samples from birds confiscated from illegal handlers. http://dx.doi.org/10.17504/protocols.io.bac3iayn. [PROTOCOL DOI].

### Genetic analysis

In order to understand the genetic structure and diversity of *P. coronata* and *S. similis*, we compared individuals from the Pampa region, in which both species occur in relative abundance, with *P. coronata* individuals from the Chaco/Pantanal region and *S. similis* from the Atlantic Forest region. There were few samples of *P. coronata* from the Atlantic Forest and of *S. similis* from the Chaco/Pantanal because these are not major occurrence regions for these species. This strategy also allowed us to compare two regions of more similar vegetation patterns (Pampa vs. Chaco/Pantanal, in the case of *P. coronata*) with two regions with more different vegetation patterns (Pampa vs. Atlantic Forest, in the case of *S. similis*). Further information about the genetic sampling is given in S2 Table.

DNA was extracted from the blood in FTA cards using the PureLink Genomic DNA Mini Kit (Invitrogen), and the fragment of the mitochondrial gene NADH dehydrogenase subunit 2 (*ND2*) was amplified using the same PCR protocol for both species. The reaction was performed with concentrations of 20 ng/µl of DNA, 1x PCR Buffer (Invitrogen), 3.5 mM MgCl2, 0.2 mM dNTPs, 0.2 pmol/µl of each primer, and 0.04 U/µl of Taq Platinum DNA Polymerase (Invitrogen). The primers used were MetL (described by Hailer et al. [31], sequence 5’-AAGCTATCGGGCCCATACCCG-3’) and RND2A (this study, sequence 5’-CCTGAGTTGCATTYAGGGG-3’), and the PCR conditions were as follows: 94°C for 2 min, 35 cycles of 94°C for 30 s, 59°C for 30 s, 72°C for 60 s, and a final extension of 72°C for 8 min. The amplification was confirmed through electrophoresis in a 1% agarose gel. The amplified products purified enzymatically with exonuclease I (GE Healthcare) and Sanger sequenced by ACTGene Inc., Brazil.

### Data analysis

For field work data, descriptive variables for each capture were recorded in a spreadsheet (S1 Table). A principal component analysis was performed based on Gower’s universal similarity index using Past v.3 (University of Oslo, Oslo, Norway. https://folk.uio.no/ohammer/past/). We used the Spearman correlation coefficient to relate the principal component with each of the studied variables. In addition, the association among categorical variables was studied using the Pearson chi-square test for association. These tests were conducted in Minitab v. 18 (State College, Pennsylvania, USA. http://www.minitab.com) using 0.05 as the significance threshold. Responses were also accessed using descriptive statistics.

For molecular data, DNA sequences were assembled and aligned in the software Geneious v.10.2.3 (https://www.geneious.com) and checked by eye in MEGA X v.10.0.0 [32]. We used DnaSP v.6.12.03 [33] to define all different haplotypes whose evolutionary relationships were represented using a Median-joining network [34], as estimated in the software PopART (http://www.popart.otago.ac.nz). Standard genetic diversity indices, including Tajima’s D [35] and Fu’s FS [36] neutrality tests were estimated using the program Arlequin v.3.5.2.2 [37]. We also used this program to quantify the level of genetic structure using both hierarchical and non-hierarchical Analysis of Molecular Variance (AMOVA) [38]. The hierarchical AMOVA was performed for each species using the three collection regions (Pampa, Atlantic Forest, and Chaco/Pantanal). Non-hierarchical AMOVA was performed only for the Pampa region to explicitly compare the level of structure in the same region using a comparable sample strategy for both species.

Finally, to understand the past demography of both species, Bayesian Skylines plots (BSP) [39] were generated for the total population and the main occurrence regions of each one of them, using BEAST v.2.6.1 [40]. We used 10,000,000 MCMC steps, sampling every 1,000 steps and discarding the initial 10% of the sampling as burnin. We used a partition scheme allowing each codon position to have a different substitution model, which were estimated in MEGA X v.10 [32]. We checked sampling sufficiency of the MCMC and built the BSP in Tracer v.1.7.1 [41]. The molecular substitution rate for the ND2 gene was calculated as described previously [42], assuming the “calibration set 2” for a 45g bird [43,44].

## Results

### Capturing birds in the wild

In total, we performed 223 captures: 122 Saltators and 101 Cardinals (S1 Table). Net-trapdoors accounted for 36% of the captures, of which 97% were *S. similis* (Fig. 3). Catches using the loop method were 33% of the total, with a high prevalence of *P. coronata* (89%). The mist nets provided comparable catches in both species under study, with 53% being *S. similis*.

**Fig. 3.**
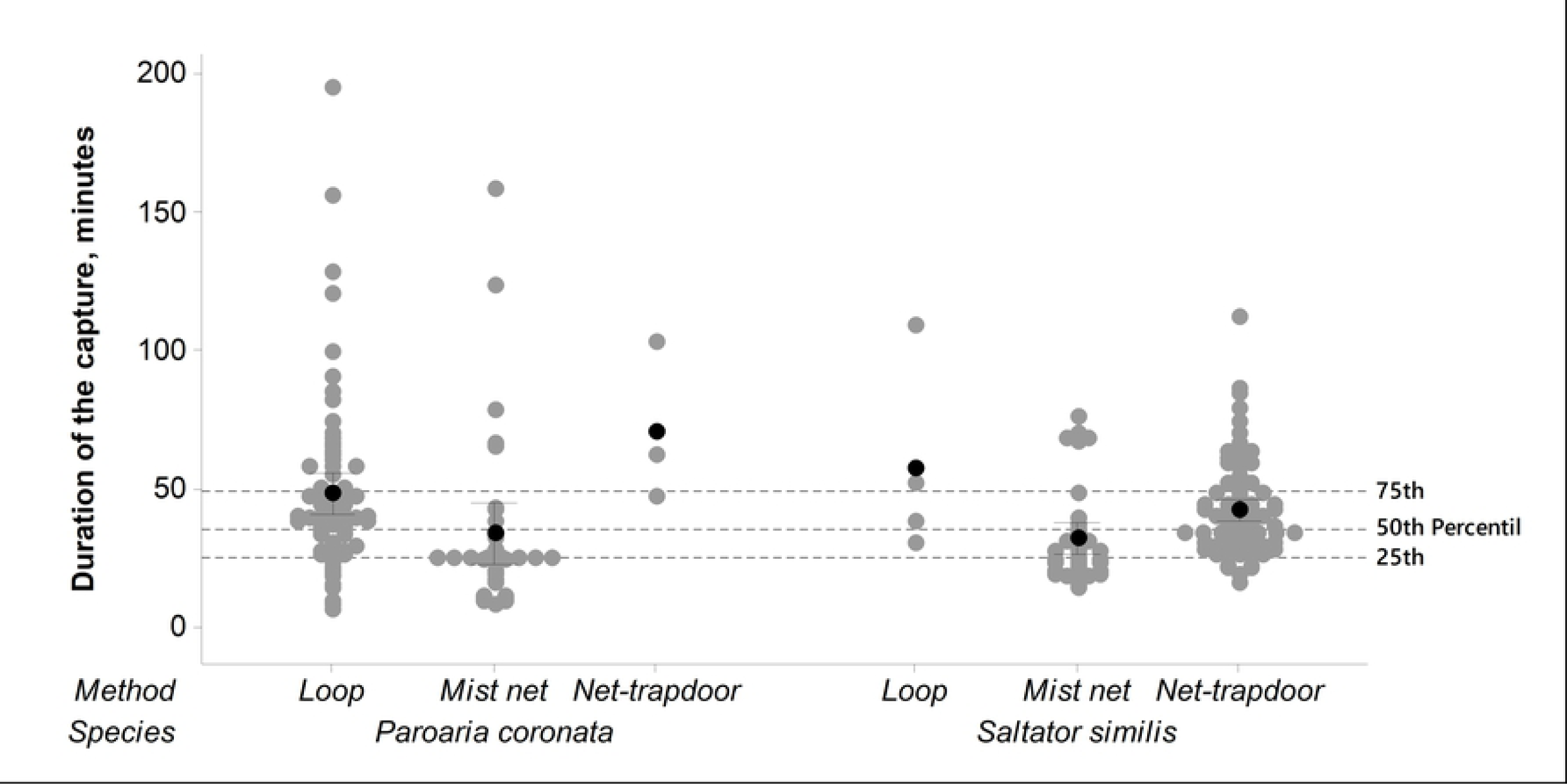
Frequency and duration of each capture according to bird species and capturing method.

On average, it took 41.7 minutes (± 1.69 minutes) for each bird capture, with similar values between species. Catches using the mist net were faster (32.8 ± 2.9 minutes) compared to captures performed with the net-trapdoor (43.1 ± 1.9 minutes) or loop (48.5 ± 3.6 minutes) methods. The mist net also accounted for 62% of the captures performed in less than 25 minutes, which correspond to the first quartile of the database. The principal component analyses show that capture method was the most important factor discriminating between *P. coronata* and *S. similis* captures (Fig. 4).

**Fig. 4.**
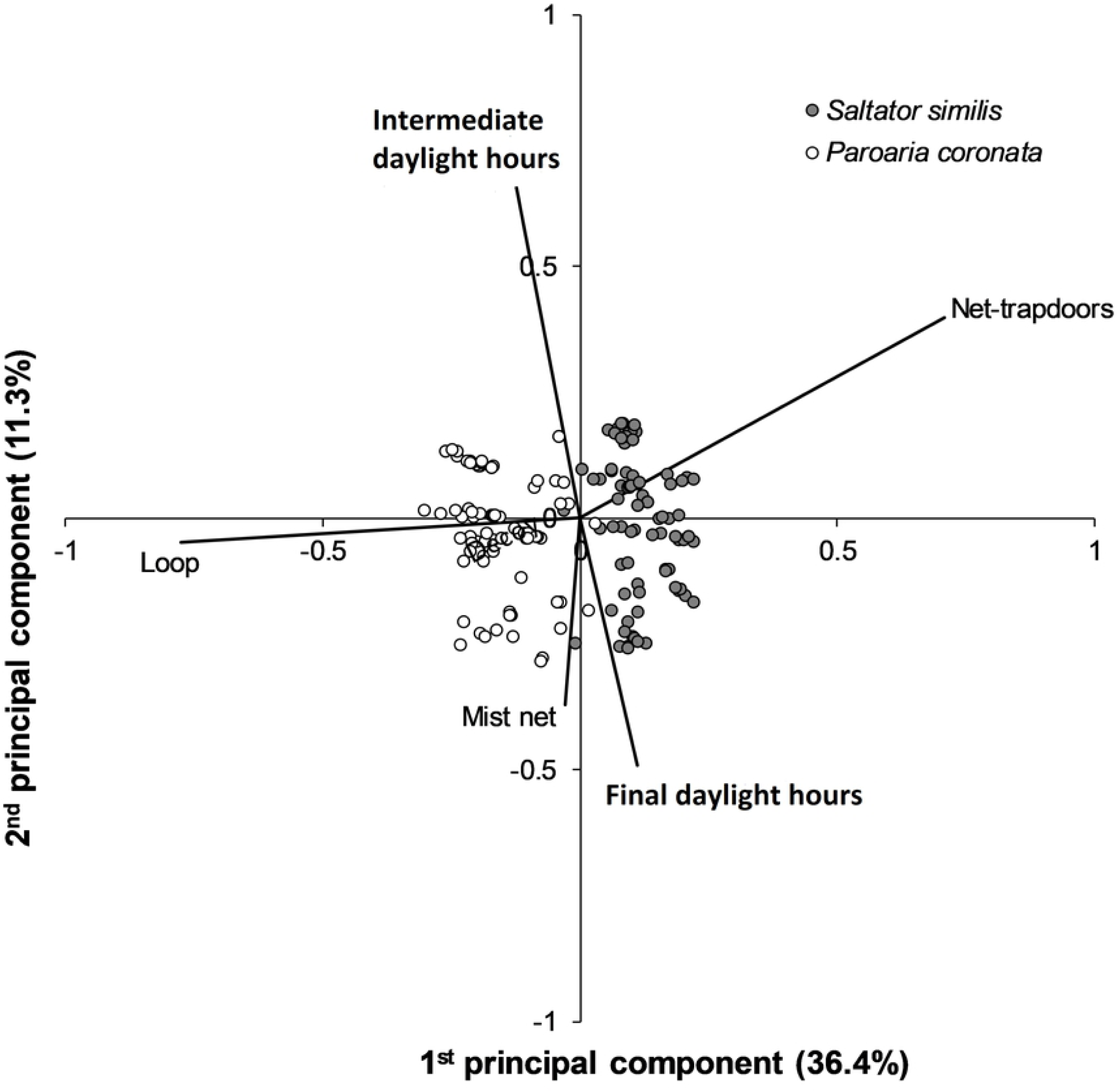
Principal component analysis indicating captures (dots symbols) and the variables having the highest explanatory power (lines).

Characteristics of the decoy were associated to all studied variables in the captured birds (Table 1), showing an important effect of the decoy in the capturing process. Some of these associations were very clear even before the statistical analysis. For instance, all captures involved conspecific decoys. Male decoys accounted for 99% of the captures of *S. similis*, while the captures of *P. coronata* were well distributed among decoy sexes (couple: 42%, male: 33%, and female: 25%). Among the female birds sampled, only 26% were captured using female decoys. A total of 60% of the female birds were captured using loop (almost exclusively *P. coronata*), while 75% of the male birds were captured using net-trapdoor or mist net, even though the low capture of female and young Saltators may bias these results. Decoys with a high ranking for dueling disposition and vocal performance (S3 Table) accounted for 96 and 98% of the fastest captures (performed in less than 25 minutes), respectively. Seven birds (∼3%) died due to capturing/sampling procedures.

**Table 1.** Probability of association between characteristics of the captured bird (sex, species, and age—or duration of the capture) with characteristics of the decoy or with the method applied in the catch

| Decoy / Method | Captured bird / Duration of capture |  |  |  |
| --- | --- | --- | --- | --- |
|  | Sex | Species | Age | Duration<sup>2</sup> |
| Sex | **<0.001** | **<0.001** | **0.001** | **0.024** |
| Species | **<0.001** | **<0.001** | **0.009** | **<0.001** |
| Dueling disposition<sup>3</sup> | **<0.001** | **0.047** | **0.015** | **0.037** |
| Vocal. performance<sup>3</sup> | **0.027** | **<0.001** | **0.005** | **0.045** |
| Method | **<0.001** | **<0.001** | 0.095 | **<0.001** |

Probabilities obtained in the Pearson chi-square test for association. Significant values are shown in bold. <sup>2</sup> Duration was ranked in four classes according to the quartiles. <sup>3</sup> Each decoy was ranked in three classes according to its behavior and vocalizations during captures (S3 Table).

### Sanitary tests

Data on the PNSA-associated tests performed on samples from both free-ranging and confiscated *P. coronata* and *S. similis* are presented in S4 Table. The serological study revealed no presence of antibodies against Newcastle disease, irrespective of the origin of the bird. *Salmonella* spp. isolation resulted in only one positive sample: *S. enterica* serovar Cerro from a free-ranging *P. coronata*. Two tests were applied for the detection of *M. gallisepticum* (MG). The RPA test indicated positive results for 25% of the samples from free-ranging birds, but none was confirmed by the PCR assay. On the other hand, seized birds showed positive results for MG in the RPA test in 53% of the samples, which was reduced to 13% when samples were tested by PCR. Most disagreement between the tests were ‘false-positive’ results. Despite this variation, results obtained by the RPA method were highly correlated (*P* <0.001) with the results obtained by PCR when assessed by a Pearson chi-square test.

### Genetic structure and diversity

DNA sequence analysis resulted in an alignment of 977 bp and 810 bp for *P. coronata* and *S. similis*, respectively. All new sequences were deposited in GenBank (S2 Table). Genetic diversity indices for both species in all sampled regions are shown in Table 2. Overall, there was higher genetic diversity in *S. similis* than in *P. coronata* for both the whole sample as well as in the Pampa region. However, we found contrasting patterns between species when the two major regions of occurrence were compared. While for *S. similis*, the Pampa and Atlantic Forest have comparable levels of diversity, for *P. coronata*, the Pampa has only a fraction of the diversity contained in the Chaco/Pantanal region, which is also clear from the haplotype network (Fig. 5). For both species, the sample size in regions of minor occurrence (Atlantic Forest in the case of *P. coronata* or Chaco/Pantanal in the case of *S. similis*) is too low to allow for a through characterization of the genetic diversity.

**Table 2.**
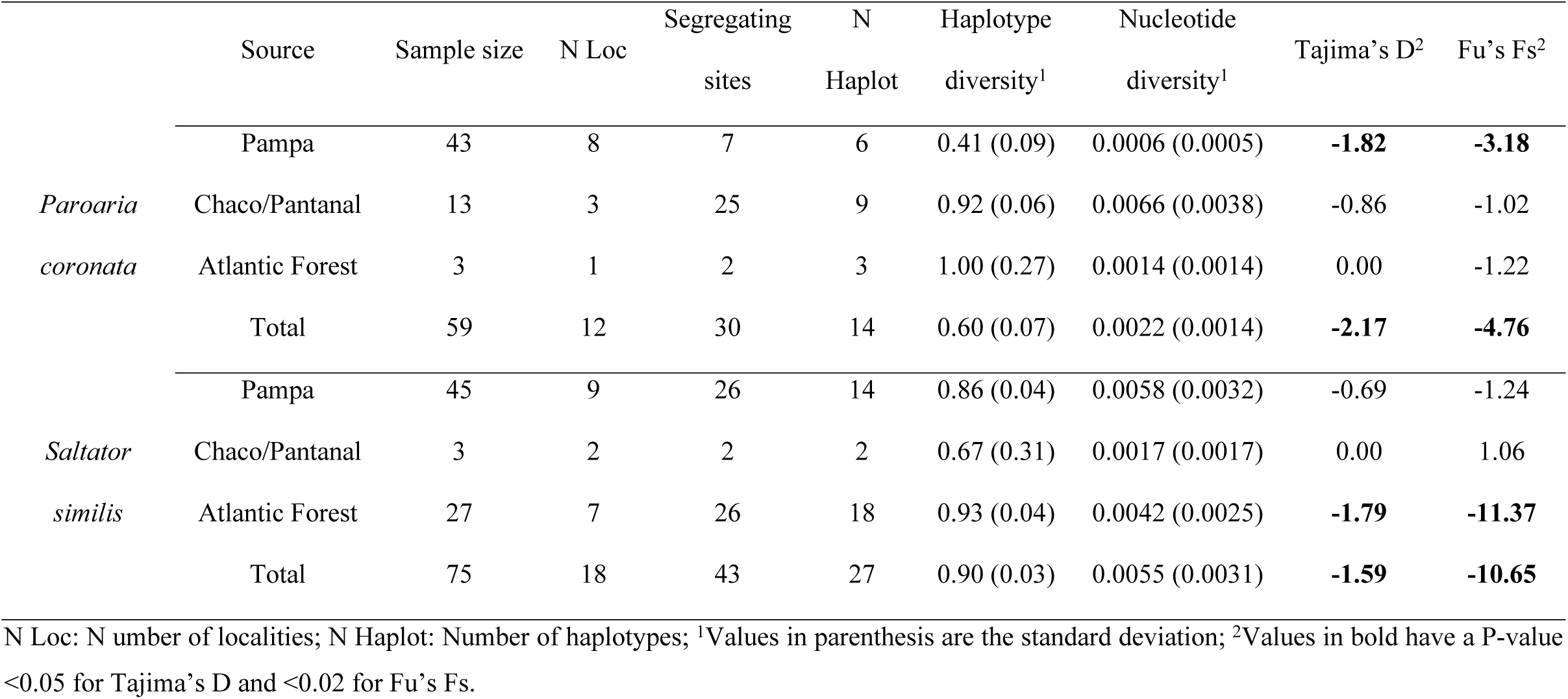
Standard genetic diversity indices for *P. coronata* and *S. similis.*

**Fig. 5.**
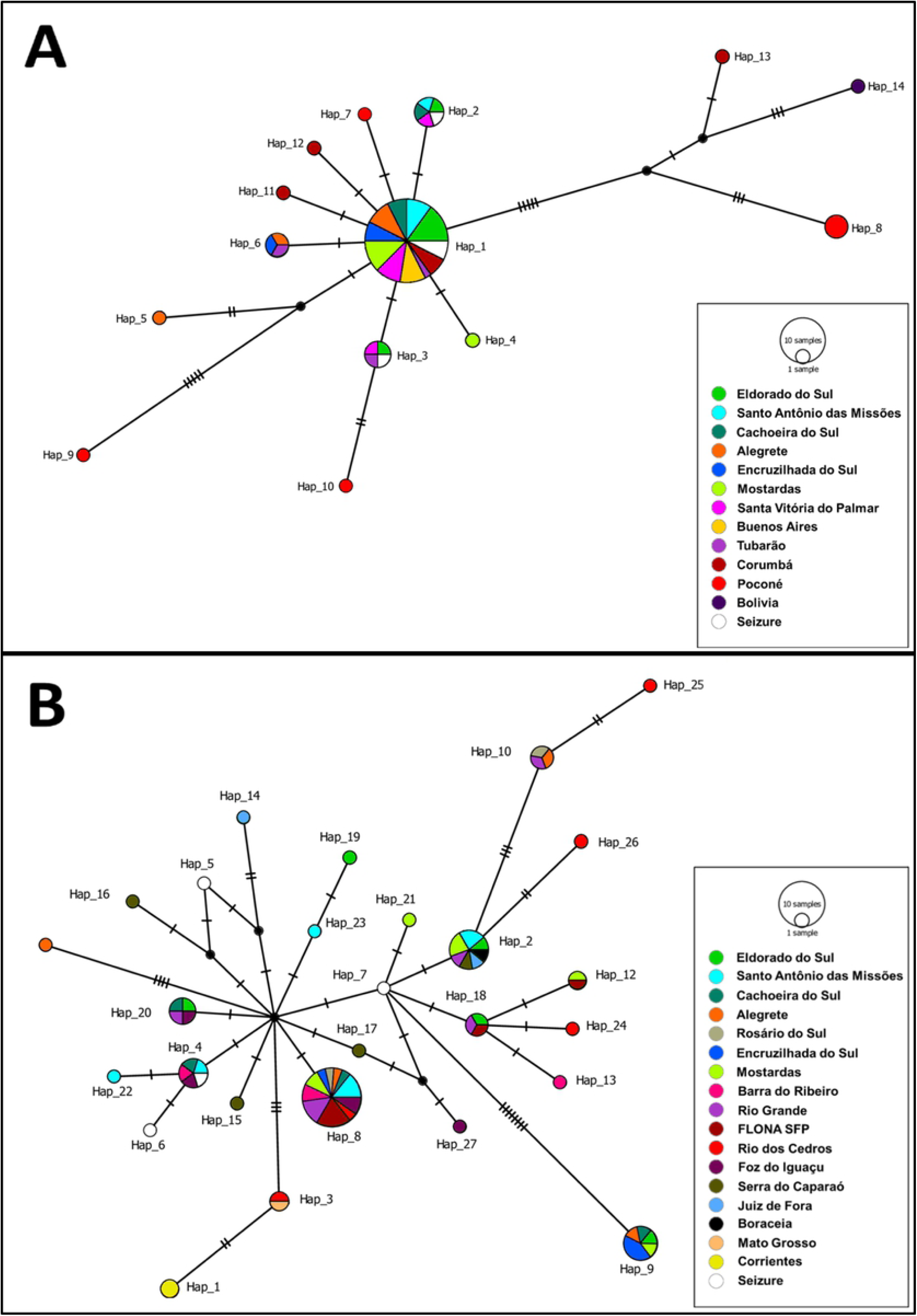
Median-joining network for the haplotypes of *P. coronata* (A) and *S. similis* (B). The circle’s size is proportional to the number of individuals in the sample with that haplotype. The transversal lines in the connectors represent the mutational steps.

The haplotype network for both species (Fig. 5) also suggests a relatively low degree of genetic structure among regions for both species. Indeed, when all regions are considered, 26.18% of the total genetic variance (*Φ*_*CT*_=0.2618; *P*=0.040) occurs among regions for *P. coronata*, while for *S. similis*, this value is only 12.94% (*Φ*_*CT*_=0.1294; *P*<0.001). On the other hand, most of the genetic variation occurs within populations: 65.17% (*Φ*_*ST*_=0.3483; *P*<0.001) and 82.72% (*Φ*_*ST*_=0.1728; *P*=0.011) for *P. coronata* and *S. similis*, respectively. The genetic structure among populations within regions corresponds to the remaining portion of genetic variance, but this value was not significant for both species (*Φ*_*SC*_=0.1171; *P*=0.069 and *Φ*_*SC*_=0.0499; *P*=0.090 for *P. coronata* and *S. similis*, respectively). When only the Pampa region is considered, there is no significant genetic structure for either species (*Φ*_*ST*_=-0.0510; *P*=0.923 and *Φ*_*ST*_=0.0711; *P*=0.107 for *P. coronata* and *S. similis*, respectively), corroborating the high general genetic homogeneity within regions shown in the previous analysis.

We found evidence for population growth in both species judging by both Tajima’s D and Fu’s FS statistics (Table 2). However, while in *P. coronata* this signal is only found in the Pampa region, in *S. similis*, it is exclusive from the Atlantic Forest region. The Bayesian Skyline analyses are congruent with these signals (Fig. 6). Population expansion in *P. coronata* seems to be more recent (∼50,000 years ago) and less dramatic than in *S. similis*, whose expansion seems to have occurred ∼350,000 years ago.

**Fig. 6.**
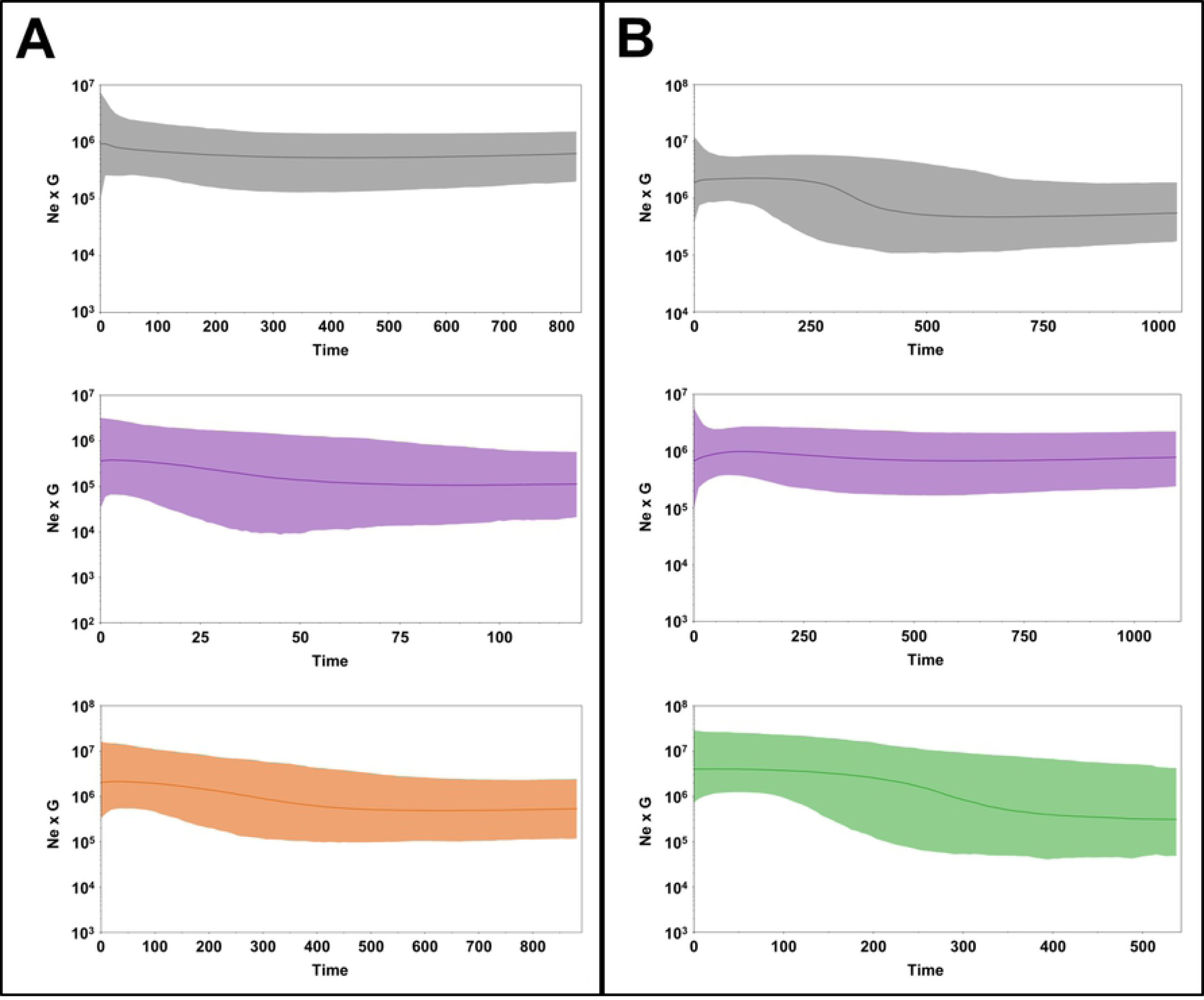
Bayesian Skylines plots for *P. coronata* (A) and *S. similis* (B). The time scale is in thousands of years. The effective population size is multiplied by the generation time in the y-axis (Ne x G). The colors indicate the region of the population considered: Total population—grey; Pampa—purple; Chaco/Pantanal—orange; Atlantic Forest—green.

## Discussion

Our planet faces many biodiversity conservation challenges to fulfill the needs of an ever-growing human overpopulation [3,4]. Considering the limitation of financial resources for conservation, it is understandable that non-threatened species do not retain high conservation priority [3,5,6]. Indeed, neither the IUCN [5,6] nor the Brazilian Ornithological Society [7] recommend translocation as a valuable conservation tool for non-threatened confiscated wild birds, as the cost of returning birds to the wild in a responsible manner can be prohibitive [6]. However, options other than humane killing of several thousand wild passerines deserve consistent investigation, since euthanizing a high numbers of wild birds annually [45,46] involves ethical issues and may be incongruent with species conservation, especially in the long-term [1,3]. In fact, there has been some evidence that common species also can be susceptible to population declines [47]. The IUCN recognizes that returning confiscated animals of low conservation value to the wild may be valid, given that there are available resources and that translocation is undertaken in accordance with standard conservation guidelines, which should be based on genetic and sanitary information in addition to a number of other studies [5,6]. *S. similis* and *P. coronata* are among the most trafficked passerine species in Brazil [45,46,48,49], which configures a legal and conservation challenge even if these species are considered as of least concern on the IUCN red list [18]. Thus, genetic [50,51] and health [52] data from these species may help to substantiate decisions on rehabilitation/release programs.

### Capturing and sampling wild birds

In this study, after consecutive unsuccessful attempts at capturing live birds, we used live decoys in association with cages mounted with net-trapdoors, Indochinese loops, and mist-nets. Live conspecific decoys have long been among the poachers’ practices for supplying illegal trade with wild birds [17]. This method simulates a territorial intrusion, which provides adequate capture opportunities, especially during the breeding season, since birds are prone to fight for defending their territories and are willing to mate [53]. Previously, baiting an area for a couple weeks to entice birds could facilitate the captures [12]; however, most of the capturing areas were visited only at the capture time.

Of the 223 catches, 83 were trapped in the net-trapdoors, of which almost all were *similis*, which readily engage in combat. On their turn, *P. coronata* are shyer, evaluating their opponents for long periods, hardly perching on unknown devices, and avoiding shaded areas that are suitable for mist netting. In this context, the use of loops camouflaged under a perch allowed the capture of this elusive species, even with the highest mean waiting-time recorded in the study. Loops were used to capture *S. similis* only in specific occasions, such as when it systematically perched in the same place and did not attack the decoy’s cage. Mist nets provided comparable catches in both species under study (Fig. 3, Fig. 4).

The average target-catches per day was ∼ 2.5, much lower than the 10.2 reported recently [14]. Unlike the aforementioned study, we spent considerable time on the road. Furthermore, the increase in the number of daily captures over time (S1 Table) suggests the importance of team learning by practice [54,55]. For example, mist-netting masked by medium height vegetation or wood board fencing, under cloudy or shaded conditions, created additional opportunities for capturing open area birds such as *P. coronata*. Although with a greater effort for installing and linked to most of the non-target captures, the mist net was the fastest (33 minutes) capturing method for both species (Fig. 3).

During the three-year period, only four out of 24 birds that were selected as live decoys showed persistently high dueling disposition. The head down and retracted tuft of mute Cardinals and the ruffled feathers, low chirping, and open fallen wings of Saltators were submission behaviors observed under the free-ranging rival challenge. It has also been shown that in some bird species, males utilize vocal performance to evaluate competitors [53,56]. In our study, one specific *S. similis* decoy that whistled an obvious longer and louder terminal trill was associated with many catches (n=58). This individual likely elicited a wider territorial response to the song performance of this “intruder”, similarly to what has been reported for *Zonotrichia leucophrys* males [56]. Thus, the efficiency of the decoy and, consequently, of the capture was associated with the level of the decoy’s song performance or underlying health and vigor [53]. In most occasions in which decoys became mute and submissive, the free-living conspecific bird gave up the fight and left. However, as long as we had a replacement decoy, methods were effective and the target species were not captured, mostly when we couldn’t find them.

Data on mortality rates in surveys involving bird capture are scarce and usually rely only on mist-netting [57,58]. While these studies report much lower mortality rates, they do not involve sampling of biological tissues. It is likely that most of the birds that died during collection, suffered from hemorrhages secondary to blood sampling. As an attempt to minimize such losses (S1 Table), blood collection was limited to 0.1 ml per bird, instead of 0.2 ml as we did at the beginning of the fieldwork.

### Sanitary and health issues

While there is a long list of diseases affecting wild birds [59], we decided to investigate only diseases addressed on the PNSA (i.e., Salmonellosis, mycoplasmosis, and Newcastle disease) due to its associated economical relevance [60] and because investigations about PNSA-associated pathogenic agents in wild passerines are rare.

All samples tested for the presence of anti-NDV antibodies were negative in this study. Comparable results have been divulgated in NDV serological surveys in seized Passeriformes, even though positive results have been observed in other avian orders [61]. Monitoring free-ranging wild birds have shown the same tendency [62].

*Salmonella Enteritidis* and *Salmonella Typhimurium* are, respectively, the serotypes most prevalent and most commonly associated with disease in wild birds [63]. However, prevalence of *Salmonella* in wild birds seems to be low and expositions have been attributed to man/domestic animals-contaminated environments [64]. Indeed, previous studies estimated the prevalence of *Salmonella* in samples of wild birds, including in passerines confiscated from illegal traffic, at between 1–7% [59,65–67]. In our sample, only one fecal sample (0.7%) was positive for *Salmonella* cultivation and isolation, which is consistent with a previous study on *Paroaria* sp. in which *Escherichia coli* and *Klebsiella* spp., among others, were detected in much higher numbers than *Salmonella* [67]. The positive sample was *Salmonella enterica* serovar Cerro (Group K). This serovar has been linked to poultry feedstock and rations [68] and commercial laying farms [69] and has high prevalence among cattle herds [70], which was the case here, even though the meaning of this finding for the health of this free-ranging Red-crested Cardinal population remains to be investigated.

We found higher prevalence for MG using the RPA test compared to the qPCR assay for both free-living and seized birds, which was expected due to the limited accuracy of RPA [29]. MG-prevalence in poultry and wild birds has usually been higher under RPA than in other tests [71–73]. The qPCR-based estimate of 13% in MG prevalence in the seized group is comparable to what has been found in other avian hosts [74–76]. Similarly, the higher MG prevalence in seized birds probably reflects the poor hygiene and stressful conditions to which these birds are subjected [59]. Even though MG exposure has been reported for wild birds [71,74,77], the lack of qPCR-based positive samples for MG is in agreement with another Brazilian study that sampled wild birds in poultry farms surrounded by forest remnants [78]. Even though none of the positive individuals in our sample showed any clinical evidence of infection, differential MG-susceptibility across bird species has been observed after experimental infections that resulted in both clinical and subclinical symptoms [79], which also occurred in wild birds [74,77,80]. Disease outbreaks in free-ranging and captive wild bird have been recorded, suggesting a potential reservoir for the pathogen in nature [81–83], reinforcing the need of a reliable MG assay for elucidating the role of MG in populations of wild birds.

### Genetic diversity, structure, and implications of outbreeding depression

The two species revealed different levels of genetic diversity and different evolutionary demographic histories. *S. similis* had overall higher genetic diversity, a more ancient population expansion restricted to the populations in the Atlantic Forest (while the Pampa population remained constant), and similar levels of diversity between the Pampa and the Atlantic Forest, its major regions of occurrence. On its turn, *P. coronata* had lower diversity, a weaker and more recent population restricted to the populations in the Pampa (while the Chaco/Pantanal population remained constant), and lower diversity in the Pampa compared to the Chaco/Pantanal region (Table 2, Fig. 5, Fig. 6).

When compared to other passerines, mainly from open areas and/or dry forests in Southern South America [84,85], *P. coronata* from the Chaco/Pantanal had a high diversity at both the haplotype and nucleotide level. Even though some population of *Gubernatrix cristata*, a highly endangered Thraupidae, had higher nucleotide diversity values [84], this study used the mtDNA control region (mtDNA-CR) as a genetic marker, which evolves faster than ND2 [86]. On the other hand, *P. coronata* from the Pampa had a low diversity at both indices. This may indicate that the population inhabiting the Pampa biome is a recent offshoot from the Chaco/Pantanal region, which may have acted as a large source population for this species. This is compatible with the exclusive signal of population expansion in the Pampa around 50,000 years ago (Fig. 6), as population growth is usually expected in refuge areas [87].

Compared to other passerines more associated with the Brazilian Atlantic Forest and other associated humid forests, *S. similis* revealed a high diversity both in the Pampa as well as in the Atlantic Forest regions. Considering studies that used mtDNA coding genes only [88–95], *S. similis* from the Atlantic Forest had the highest value for a forest-associated species (only lower than *P. coronata* from the Chaco/Pantanal), while the Pampa population had the fifth highest value. Haplotype diversity values for both populations are also among the highest for passerines associated with the Atlantic Forest [88–95]. The exclusive signal of an ancient population expansion for *S. similis* in the Atlantic Forest region but not in the Pampa is curious. First, most studies that detected a population expansion in specific populations or phylogroups suggest much more recent timescales (similar to *P. coronata*) [90,92,94,96]. The two exceptions that report older expansions are *Schiffornis virescens* [91] and *Basileuterus leucoblepharus* [92], around 150,000–300,000 years ago. The high genetic diversity and lack of a signal of population expansion in the Pampa is intriguing. It is unlikely that the Pampa behaved as a refuge area for this species, which is strongly associated with humid forests northern to the Pampa region. Besides, the Atlantic Forest has higher genetic diversity than the Pampa, which is not compatible with a scenario in which the population in a refuge – the Pampa – occupies the Atlantic Forest and expands thereafter. Taken together, the lack of strong genetic structure between these populations may indicate that the Pampa was occupied in consequence of the population expansion in the Atlantic Forest but subsequently lost genetic diversity, which eroded the signal of an ancient expansion.

We also found a very weak genetic structure for both *P. coronata* and *S. similis* (Fig. 6). In the Pampa and associated dry forests, both *Gubernatrix cristata* [84] and *Phytotoma rutila* [85] showed stronger structure among habitats (*Φ*_*ST*_∼0.45). For forest species, most studies reported the presence of strongly differentiated phylogroups either between different habitats or across the Atlantic Forest [88–90,93–99]. Again, the exceptions are *Schiffornis virescens* [91] and *Basileuterus leucoblepharus* [92], which showed no phylogeographic breaks across its distribution in the Atlantic Forest. The lack of genetic structure in *S. similis* is corroborated by a study that, based on nine microsatellite (SSR) loci, reported that only 0.1% of the total genetic variation occurred among populations distributed in different Brazilian biomes (Atlantic Forest, Cerrado, Caatinga and ecotones between these) [50]. Similarly, even though no study has characterized *P. coronata* populations for SSR variation, the closely related species *P. dominicana*, which occurs in the Brazilian northeast, showed that only 3% of the total genetic variation occurred among populations. [50]. Thus, the high genetic connectivity among populations does not seem to be an artifact from using only mtDNA.

The shallow population differentiation found in this study may have important consequences for conservation policies. Even though genetic distance is not a good predictor of outbreeding depression [100], outbreeding depression requires different genetic adaptations to local environments that will be broken by crossing individuals from exogenous populations [9,10,101]. The high historic genetic connectivity across populations of both species suggests that the overall risk of outbreeding depression is low [101].

### Concluding remarks

The management of birds seized from illegal traffic is a complex and difficult question; Even though the recommendation for euthanizing individuals belonging to species of low conservation value may be justified [6,7], could new information for focal species reduce the perceived risk of translocation to the point that this could become a standard practice? In this study, we showed that it may be feasible to screen seized birds for several pathogens and that while the overall prevalence rates are low, the risk of disease transmission during bird management or after release in natural environments justifies a systematic sanitary screening. In addition, applying an adequate quarantine period for all seized birds seems to be both feasible and indispensable for maintaining proper sanitary conditions of the wild bird populations in translocation/reintroduction initiatives. Furthermore, systematic pathological investigations on confiscated birds dying during the quarantine/rehabilitation processes may point to additional relevant conditions to be included on the health screening.

The shallow genetic structure found for both species suggest that the overall risk of outbreeding depression following translocation is low. Yet, additional criteria could be used to choose an area for translocating individuals. For example, using different regional vocal “dialects” as a proxy for the original parental population of an individual [102]. Vocal plasticity exhibited by several species may indicate that, for the undetermined cases, reintroduced birds will be able to adapt to their release sites [102]. Another eventual strategy could be releasing the birds close to the apprehension site. Conservationist policymakers should consider when translocating seized individuals into wild populations could represent a safe and useful strategy for maintaining large and healthy wild populations for species highly targeted for illegal trade, irrespective of their conservation status.

## Acknowledgements

We are grateful to CMPC Celulose Riograndense Ltda. for financial support and to the land owners who allowed access to their property to conduct fieldwork, especially to Biesdorf Family (Eldorado do Sul), Helena and Edegardo Velho, Ronaldo Bicca, André Carvalho (Mostardas), Paulo and Fernando Menezes (Cachoeira do Sul), Ione Ferreira, César Silveira (Encruzilhada do Sul), Teófilo Garcia (Santo Antônio das Missões), Ivan Pereira, Arcídio Trojan (Alegrete), Lívia Quoos, Mauro Amaral (Rosário do Sul), Ênio Brito (Santa Vitória do Palmar), and Gildomar Monte (Rio Grande), but also to André Thuronyi, Guilherme Arruda, Denise Senna (Poconé), Débora Murgol (Santo Antônio do Pinhal) and João Venturini (Corumbá). We thank Armando Garcia, Marcelo Maronna, Cristina Trein, and João Almeida for connecting us with land owners. We are also grateful to ICMBio and IBAMA for providing us the study licenses, and to the managers of the conservation areas Edenice Souza (Floresta Nacional de São Francisco de Paula), and Soraya Ribeiro (Parque Natural Morro do Osso). We also thank Fabiana Quoos Mayer for technical assistance and Raquel Hohendorf, Diógenes Machado, Miguel Corrêa, and Nicole Böelter for their valuable help. Finally, we thank Fundação de Apoio da Universidade Federal do Rio Grande do Sul for administrative support

## Supporting information captions

**S1 Table –** Captures.

**S2 Table –** Genetic samples.

**S3 Table –** Live decoys.

**S4 Table –** Health tests.

